# Structural intermediates observed only in intact *Escherichia coli* indicate a mechanism for TonB-dependent transport

**DOI:** 10.1101/2021.03.18.436049

**Authors:** Thushani D. Nilaweera, David A. Nyenhuis, David S. Cafiso

## Abstract

Outer membrane TonB-dependent transporters facilitate the uptake of trace nutrients and carbohydrates in Gram negative bacteria and are essential for pathogenic bacteria and the health of the microbiome. Despite this, their mechanism of transport is still unknown. Here, pulse EPR measurements were made in intact cells on the Escherichia coli vitamin B_12_ transporter, BtuB. Substrate binding was found to alter the C-terminal region of the core and shift an extracellular substrate binding loop 2 nm towards the periplasm; moreover, this structural transition is regulated by an ionic lock that is broken upon binding of the inner membrane protein TonB. Significantly, this structural transition is not observed when BtuB is reconstituted into phospholipid bilayers. These measurements suggest an alternative to existing models of transport, where TonB binding alone is sufficient to produce allosteric rearrangements in the transporter. They also demonstrate the importance of studying outer membrane proteins in their native environment.

## Introduction

The passive permeation of low molecular weight solutes across the outer membrane (OM) of Gram-negative bacteria is typically facilitated by porins. However, many higher molecular weight solutes and trace nutrients, including carbohydrates, iron siderophores, cobalamin, copper, and nickel, are bound and transported across the OM by a family of active transporters that are TonB-dependent (1, 2). These TonB-dependent transporters (TBDTs) derive energy from the bacterial inner membrane by interacting with TonB, a transperiplasmic protein that interacts with the inner membrane proteins ExbB and ExbD (3, 4). This family of transporters has two distinct domains: a β-barrel formed from 22 anti-parallel β-strands, and a core or hatch domain that fills the interior of the barrel. The β-strands on the extracellular surface of the protein barrel are often connected by long loops, while short turns join the strands on the periplasmic interface. TonB interacts with the transporter through a conserved motif on the N-terminal side of the core termed the Ton box (5, 6).

The mechanism of transport in TBDTs is presently poorly understood; however, the large size of most substrates and the absence of any obvious pathway for substrate permeation (7) has led to proposals that transport is mediated by a significant conformational event that involves a partial rearrangement or full removal of the core domain from the surrounding barrel (8). Although many high-resolution structures are available for TBDTs, there is no direct evidence for a major structural change within the core of TBDTs that might indicate a transport mechanism. In the *Escherichia coli* vitamin B_12_ (cobalamin) transporter, BtuB, EPR spectroscopy shows that substrate binding unfolds the Ton box at the N-terminus and extends it into the periplasm, an allosteric event that may facilitate the binding of TonB to BtuB (9, 10); however, no other significant structural changes have been observed in the core.

High-resolution crystal structures have been obtained for a C-terminal fragment of TonB in complex with BtuB and the ferrichrome transporter FhuA (5, 6). When TonB binds, the Ton box extends from the core and interacts with the β-sheets of TonB in an edge-to-edge manner. Except for the Ton box, the remainder of the core remains folded and is essentially unchanged. Because TonB binding does not alter the core of BtuB in the BtuB-TonB structure, it has been proposed that TonB alters the core by exerting a mechanical force on the transporter, and current models for transport favor a mechanism where TonB acts by pulling the Ton box thereby unfolding the core (11-13). Models involving a rotation of TonB have also been proposed (14); however, in FhuA there are 4 to 5 unstructured residues between the Ton box and core when TonB is bound, making the transfer of torque from TonB to the core unlikely (15). Pulling models have been explored using steered molecular dynamics (MD) (11) as well as single-molecule AFM pulling experiments (12), and these studies indicate that an N-terminal region of the core (up to residue 73) is preferentially unfolded to permit the movement of vitamin B_12_ into the periplasm. This work concludes that the C-terminal region of the core is static and does not unfold during transport, a result that is consistent with denaturation experiments on BtuB (16).

An important caveat to almost all the structural work on BtuB is that it has been carried out on purified or partially purified protein where the native OM environment is no longer present. Since transport in this family of transporters has never been reconstituted, it has never been established that the isolated, purified, and membrane reconstituted BtuB is capable of transport. Recently, we developed an approach to attach spin labels to either extracellular or periplasmic sites on BtuB in intact cells, thereby permitting EPR measurements to be made under conditions where the protein is known to be functional (17, 18). Preliminary measurements made on BtuB indicate that it behaves differently in the intact cell than it does in a purified reconstituted phospholipid system. For example, a substrate-dependent change in the core domain of BtuB involving SB3 is observed *in situ* but is not seen in a detergent treated OM preparation (18). Moreover, the extracellular loops of BtuB are also highly constrained in the intact cell, and substrate-induced structural changes and structural heterogeneity that is observed for BtuB in proteoliposomes is not observed for BtuB *in situ* (19).

In the present work, we perform Double Electron-Electron Resonance (DEER) on BtuB in intact cells to determine whether structural changes take place in the core domain that are associated with substrate binding and transport. Upon substrate binding, a movement of the core is observed involving sites 90 and 93 in SB3. No other movements in the core are detected. When the ionic lock is broken between site R14 on the C-terminal side of the Ton box and site D316 in the barrel, long distance components appear upon substrate binding, indicating that SB3 can assume a state where it has moved into the BtuB barrel towards the periplasmic side of the protein. Under these same conditions, other sites on the N-terminal side of the core remain static. Since this ionic interaction would normally be broken upon TonB binding, this structural transition likely represents an intermediate that occurs during transport. This result suggests a mechanism for transport that does not involve either a pulling or rotation of TonB, but rather involves allosteric changes in the C-terminal side of the core upon TonB binding. Remarkably, these substrate-induced structural changes are not observed for purified, membrane reconstituted BtuB, which may be due to the absence of lipopolysaccharide (LPS) in the reconstituted system. The importance of the native OM environment provides an explanation for why this structural transition has not been previously observed.

## Results

### Multiple sites in the core region of BtuB may be spin labeled in intact *Escherichia coli*

To investigate movements that might occur in the BtuB core region *in situ*, pairs of spin labels were placed into the extracellular region of BtuB, with one label located at an outer loop site that is known to be relatively fixed in the cell environment and a second label located at a site in the BtuB core. Previous work has demonstrated that several sites on the extracellular surface of BtuB may be spin labeled *in vivo* using site directed cysteines and a standard methanethiosulfonate reagent to produce the side chain R1 (Fig. 1a). These included multiple sites on the extracellular loops of BtuB (18-20) as well as two sites in the core region (18).

**Figure 1:**
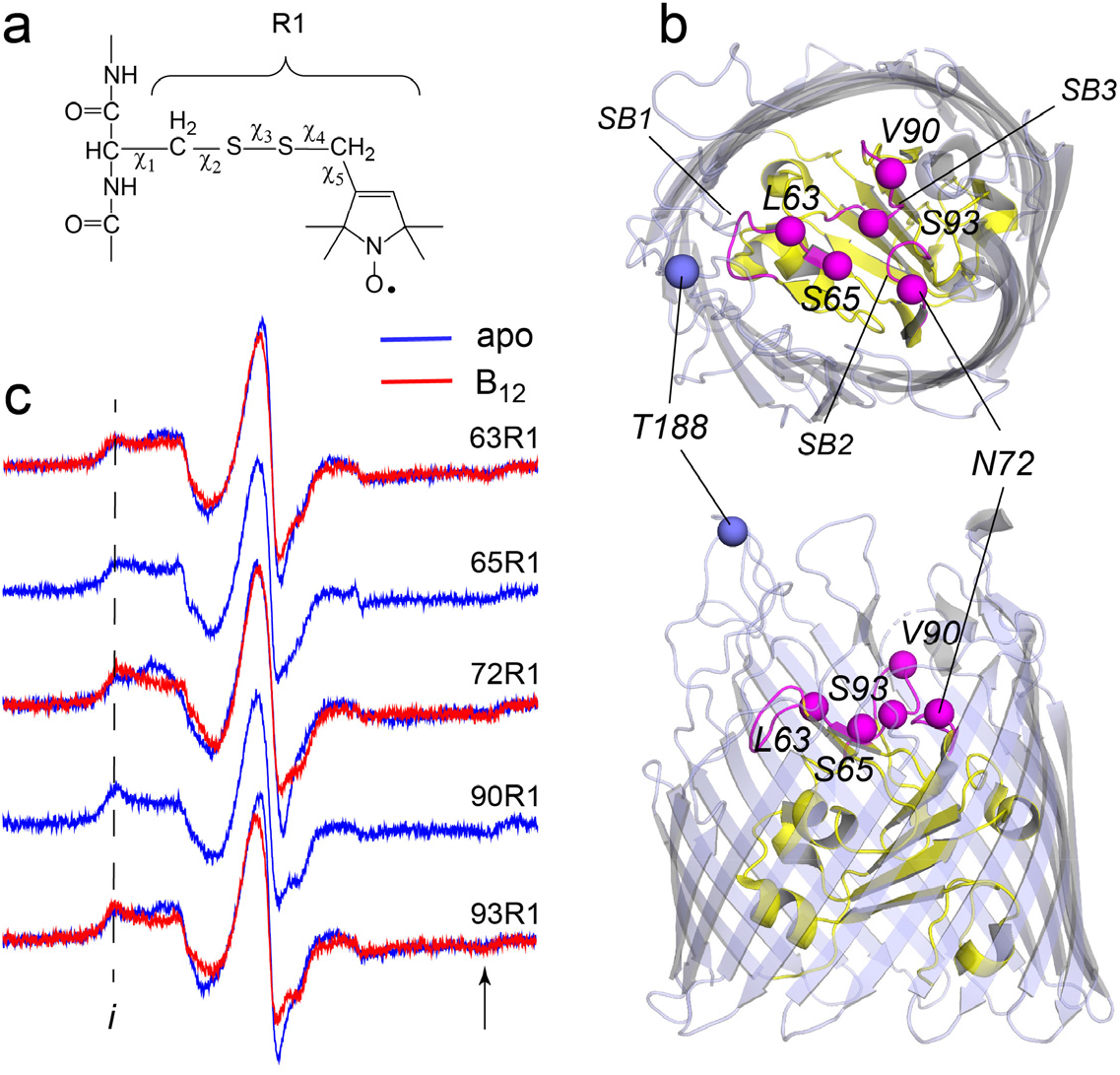
EPR spectra obtained *in-vivo* from spin labeled core sites on extracellular face of the BtuB. The spin labeled side chain (**a**) R1 was attached to sites on the extracelluar core of BtuB shown in (**b**). BtuB is shown in both extracellular (top) and side views (PDB ID: 1NHQ), with the core in yellow and barrel in light blue. The Cα atom on site 188 on the 2^nd^ extracellular loop is shown, which is used as a reference point for measurements to the core. Labeled Cα sites in the core are shown in magenta, along with substrate binding loops SB1, SB2 and SB3. In (**c**) EPR spectra are shown in the absence (apo state in blue) and presence of substrate (vitamin B_12_ bound state in red). No change in the spectrum is observed at sites 65 and 90 upon the addition of substrate. The spectra are characterized by broad hyperfine extrema (*i*) where 2Azz’ is approximately 69 Gauss, indicating an immobilized population of label. Spectra are the sum of 10 100 Gauss scans.

Recent work has also shown that the efficient incorporation of pairs of spin labels to make distance measurements using DEER required the use of a strain deficient in the disulfide bond formation (Dsb) chaperone system (18). We tested several additional single cysteine mutants in BtuB using a DsbA^-^ strain to determine whether spin labeling of additional sites in the core was possible. Shown in Fig. 1c are spectra from site 90, which was previously labeled, as wells as 4 additional sites in the core region. Sites 63 and 65 lie in substrate binding loop 1 (SB1), site 72 lies in substrate binding loop 2 (SB2), and sites 90 and 93 are positioned in substrate binding loop 3 (SB3). These spectra arise from label having more than one motional component but are dominated by a broad feature that is characteristic of a population of label with hindered motion on the ns time scale, consistent with the confined environment in the extracellular region of the core. For sites 63, 72 and 93, the addition of vitamin B_12_ alters the spectra and increases the population of the immobile component indicating that incorporation of the label at these sites has not prevented the binding of substrate. No significant changes with substrate are seen for sites 65 and 90. At site 90, substrate does bind (see below) and the lack of a change in the EPR spectrum may reflect the fact that in the apo state the label is already highly immobile. Site 65 is also highly immobile, but we cannot exclude the possibly that incorporation of R1 at this site has blocked the binding of vitamin B_12_.

### The apex of the SB3 loop in the BtuB core undergoes a substrate-dependent conformational change

For distance measurements using pulse EPR, each set of spin pairs included a label at position 188 on the 3/4 extracellular loop (the second loop connecting β-strands 3 and 4). This site was chosen as a reference point because previous work in whole cells demonstrated that this loop assumed a well-defined position and exhibited minimal or no movement upon substrate addition (19).

Shown in Fig. 2 are the results for measurements on the V90R1-T188R1 spin pair in cells. Preliminary results from this pair were presented in a previous study demonstrating the use of disulfide chaperone mutants to achieve double-labeling of BtuB in whole cells (18). The background corrected DEER data and resulting distance distributions are shown in Figs. 2b, c. Both the apo (blue) and vitamin B_12_ bound (red) distributions yield two main intramolecular peaks at 2.4 and 3.2 nm, with a substrate-dependent shift observed towards the shorter component. Predicted distance distributions were generated from the apo and vitamin B_12_ bound *in surfo* crystal structures of BtuB (PDB IDs: 1NQG and 1NQH) using the program MMM (21). The distributions generated from these structures also show a shift towards a shorter distance in the substrate bound state, which is due to the unfolding of a helical turn in the SB3 loop (22). However, the magnitude of the predicted shift is smaller than that observed by DEER. Because the position of site 188 in the 3/4 extracellular loop of BtuB is not altered with substrate addition *in-situ* (19), this structural change must involve a movement of the SB3 apex or a change in rotamers assumed by R1 at position 90.

**Figure 2:**
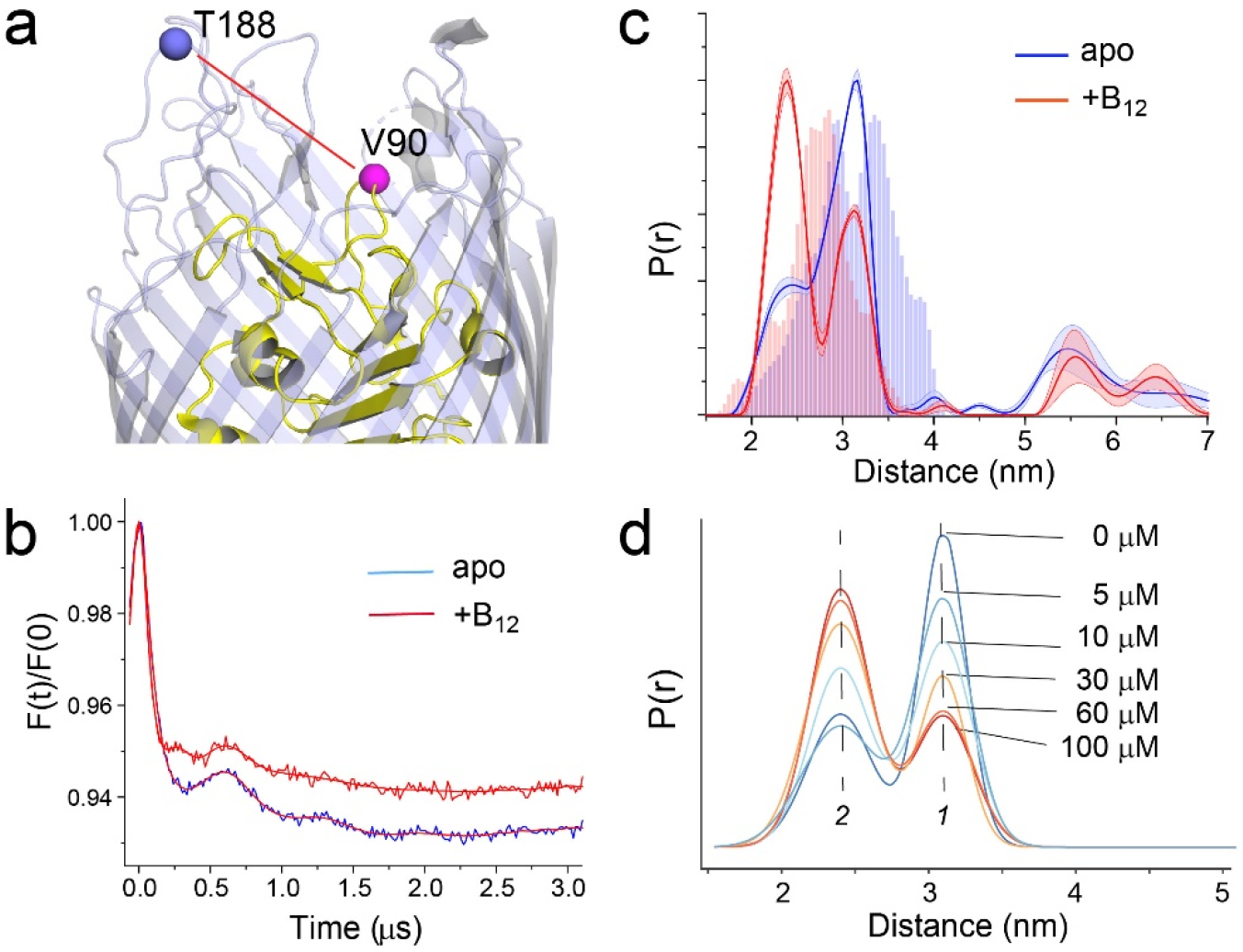
Substrate dependent shifts are detected at the apex of SB3 in whole cells. A side view of BtuB (**a**) with the locations of site 90 in substrate binding loop 3 (SB3) of the hatch domain (yellow) and site 188 near the apex of the 3/4 extracellular loop (light blue). (**b**) Background corrected DEER data for the apo (blue) and substrate bound states (red) of the V90R1-T188R1 spin pair, where the red traces represent the fits to the data. The resulting distance distributions are shown in (**c**) where the histograms represent predicted distance ranges obtained from the *in surfo* crystal structures 1NQG (blue) and 1NQH (red) using the software package MMM (21). (**d**) This structural change titrates between two states (labeled 1 and 2) with the addition of vitamin B_12_. The conversion between states saturates at concentrations above 60 µM. For the distributions shown in (c), data were analyzed using LongDistances v932 using the model-free fitting mode, whereas for distributions in (d), data were fit to a 2-Gaussian model where position, width, and amplitude were variables in the fit.

We also titrated this structural change by measuring the distance distribution with increasing concentrations of the vitamin B_12_ substrate, where the result is shown in Fig. 2d (raw and background corrected DEER data are shown in Fig. S1). For this analysis data were processed using a model-based approach with 2 Gaussian components where the position, width, and amplitude was varied. As seen in Fig. 1d, there is a strong progressive response to increases in substrate until saturation is reached in the range of 30 to 60 μM vitamin B_12_. This titration likely reflects substrate loading. Because substrate concentrations greatly exceed the affinity of vitamin B_12_ to BtuB (23), the saturation point likely reflects the concentration of BtuB in our sample, and is roughly consistent with the spin concentrations expected from the EPR signal intensity.

### Substrate dependent shifts in the extracellular face of the hatch domain are localized to the SB3 loop

To determine whether the structural change observed in the apex of SB3 is limited to this site or part of a broader conformational change across the core domain, we tested additional spin pairs on the extracellular face of the protein using the core sites shown in Fig. 1b. The spin pairs examined are shown in Fig. 3a, b and include site 93, which also lies in SB3, as well as sites 63 and 65 in SB1 and site 72 in SB2. The distance distributions that result from these spin pairs are shown in Fig. 3c, along with the V90R1-T188R1 spin pair. The raw and background corrected DEER signals are presented in Fig. S2.

**Figure 3:**
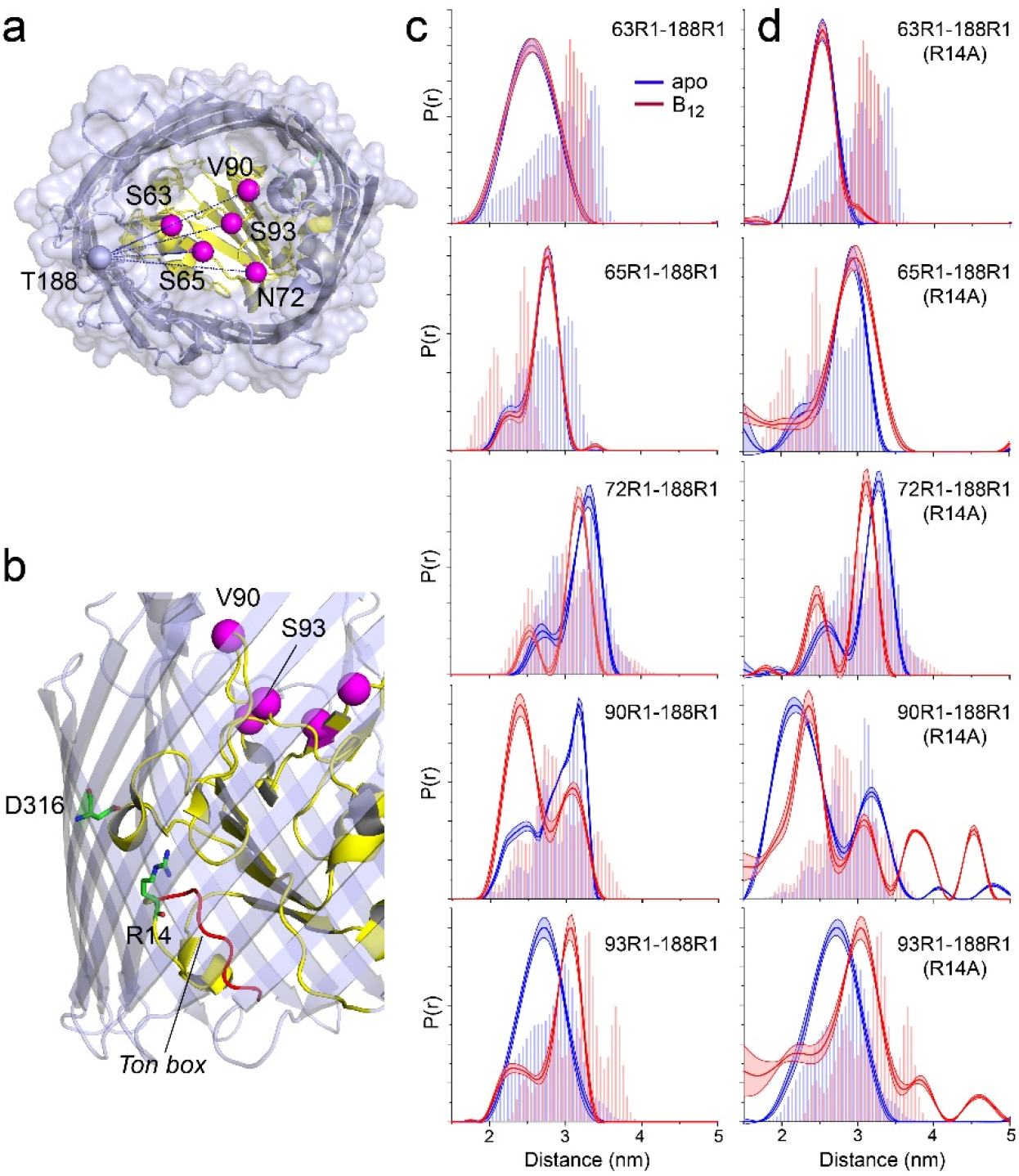
Substrate dependent conformational shifts are limited to SB3 and are altered by mutation of the R14-D316 ionic lock. (**a**) Top view of BtuB (PDB ID: 1NQH) showing the locations of the hatch sites relative to the reference site, 188, in the 3/4 extracellular loop. In (**b**) the location of the R14-D316 ionic interaction between the core and the barrel is also shown along with the Ton box (red). (**c**) Distance distributions obtained for hatch: barrel pairs in the apo state (blue) and with substrate (red). (**d**) Distance distributions obtained for hatch: barrel pairs in the apo state (blue) and with substrate (red) in the presence of the R14A mutation. Both V90R1-T188R1 and S93R1-T188R1 spin pairs show additional distances at 3.8 and 4.5 nm in the presence of R14A. Data were analyzed using LongDistances v932 and the model-free fitting regime. The shaded error bands in (c) and (d) represent variation due to background noise, start-time, dimensionality, and regularization. Histograms are predicted distances generated from the *in surfo* crystal structures PDB ID: 1NQG (light blue) and PDB ID: 1NQH (light red) using the software package MMM (21).

As seen in Fig. 3c, distance distributions for sites located outside SB3 show little evidence for any structural change upon the addition of substrate, with spin-pairs involving sites 63, 65 and 72 having nearly identical distributions for both apo and vitamin B_12_ bound conditions. This lack of a shift in these spin pairs is consistent with work showing that site 188 does not show a substrate-dependent shift in its position *in situ* (19). In the SB3 loop, however, site 93 at the edge of the loop shows a substrate-dependent change in position along with site 90 at the loop apex, although the change is much larger for site 90 (about 8 Å) than for site 93 (about 4 Å) and in an opposite direction. The structural changes measured by DEER are qualitatively consistent with the predictions from the crystal structures, which show a loss in helical structure and change in position of the loop. For the sites examined here, substrate dependent changes in the core appear to be confined to the region of the SB3 loop. Interestingly, as we demonstrate below, this substrate-dependent change does not occur when the protein is removed from its native environment.

### Breaking an ionic lock between the BtuB core and barrel triggers a large substrate-dependent change in SB3

As indicated above, BtuB is expected to bind to both substrate and TonB during transport, which will break the ionic lock between R14 in the core and D316 in the barrel (6). However, under the conditions of our experiment, BtuB is in large excess relative to TonB, perhaps by a factor of 10 to 20 or more. As a result, only a small portion of the BtuB would be bound to TonB at any time during our distance measurement.

To determine whether there might be a connection between the structure of the core and this internal ionic lock, we examined the effect of disrupting the R14-D316 interaction on the core by introducing the R14A mutation into the existing pairs of labels between site 188 in the 3/4 extracellular loop and the core. Distance measurements for the apo and vitamin B_12_ bound states made in the presence of this mutation are shown in Fig. 3d. For distance distributions involving sites 63, 65 and 72 in SB1 and SB2, the core remains largely unchanged in response to substrate and unchanged by the R14A mutation. However, distance distributions involving sites 90 and 93 in SB3 are altered by breaking the D316/R14 ionic lock.

For distances measured to sites 90 and 93, the R14A mutation has two main effects. First, for the V90R1-T188R1 pair, the substrate dependent conversion between the 2.4 and 3.2 nm distance components is absent, and the shorter distance now dominates in both apo and vitamin B_12_ bound conditions. Second, for both the V90R1-T188R1 and S93R1-T188R1 pairs, additional distance components are observed in the presence of substrate centered at 3.8 and 4.5 nm. These long components result in a distribution that is substantially broader than that predicted by the crystal structures and they indicate the formation of a novel conformation of the SB3 loop and an altered substrate binding mode.

The longer distance components that appear for the V90R1-T188R1 and S93R1-T188R1 spin pairs represent a substantial movement of the SB3 loop towards the periplasmic side of BtuB. A movement of SB3 towards the extracellular surface is highly unlikely, in part because movement in this direction would require a major unfolding of the core that we do not observe. For measurements to SB3, there were relatively few positions that were both accessible and within the range of the pulse EPR measurement, but we made measurements to site 90 from site 237, which is located near the apex of the 5/6 extracellular loop. In the apo and vitamin B_12_ bound states, the predicted distances from this site are shorter than 2 nm and are not within a range that can accurately be measured by DEER. The results are shown in Fig. S3. In the absence of the R14A mutation, no clear substrate-dependent shift in the position of SB3 is observed, which is likely due to the short distance involved. However, in the presence of the R14A mutation, a new distance component appears with substrate addition around 3 nm that is beyond the distance range predicted by the crystal structure. This is shorter than the 4.5 nm observed from position 188, which likely reflects differences in the side chain direction and the relative positions of the 3/4 and 5/6 loops.

### Substrate-dependent changes in the SB3 loop require a native environment

Two conformations are observed by crystallography for SB3. In the apo structure (PDB ID: 1NQG), SB3 has a single helical turn and shorter conformation that we speculate may be associated with the distance observed by EPR at 3.2 nm for the V90R1-T188R1 spin pair (labeled 1 in Fig. 2d). In the vitamin B_12_ bound structure (PDB ID: 1NQH), SB3 is more extended, and this state may be associated with the distance at 2.4 nm (labeled 2 in Fig. 2d). Computational work suggests that the extended state of SB3 requires the interaction of BtuB with LPS, whereas the shorter helical state of SB3 occurs in the presence of phospholipid (24), suggesting that environment, specifically LPS, may be important in controlling the configuration of SB3.

To test for an environmental effect on SB3, we reconstituted four spin pairs of BtuB into POPC proteoliposomes. These included the V90R1-T188R1 and S93R1-T188R1 spin pairs both in the absence and presence of the R14A mutation. The dipolar evolution data and distance distributions from DEER measurements on these reconstituted BtuB samples are shown in Fig. 4 (the raw time-domain data are presented in Fig. S4). In the absence of the R14A mutation, the substrate dependent conversion to the shorter distance that was seen for the V90R1-T188R1 spin pair in whole cells (Figs. 2c and Fig. 3c) is now much more limited, with only a minor shoulder appearing around 2.5 nm. For the S93R1-T188R1 spin pair, the change seen in Fig. 3c with substrate is largely absent. The 0.4 nm substrate-dependent shift to a longer distance is absent, but the small 2.2 nm distance component is still present.

**Figure 4:**
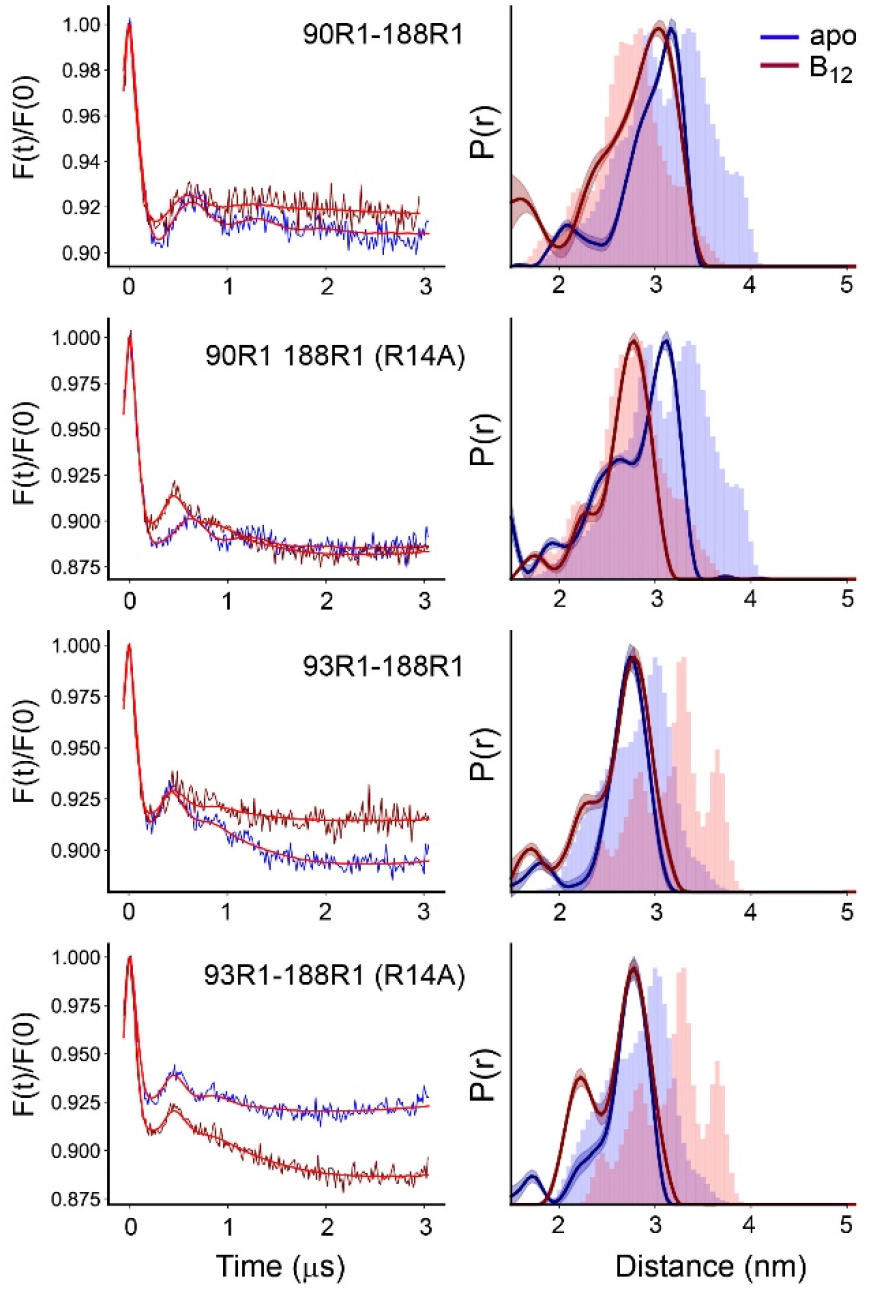
The substrate induced changes in SB3 are altered or absent in proteoliposomes. Background corrected DEER signals (left) and distance distributions (right) for the V90R1-T188R1 and S93R1-T188R1 pairs involving SB3 in the absence and presence of the R14A mutation, where the labeled BtuB have been reconstituted into POPC vesicles. Data were analyzed using LongDistances v932 and the model-free fitting mode. The shaded error bands represent variation due to background noise, start-time, dimensionality, and regularization. Histograms are predicted distances generated from the *in surfo* crystal structures 1NQG (blue) and 1NQH (pink) using the software package MMM (21).

The behavior of SB3 in the presence of the R14A mutation is also altered in the phospholipid reconstituted system. Rather than increasing the population of the shorter distance component, adding the R14A mutation to the V90R1-T188R1 pair in the reconstituted system results in a single short distance, where the resulting distribution aligns almost perfectly with the predicted distribution from the 1NQH structure. For the S93R1-T188R1 pair, R14A causes a small increase in the short distance component at 2.2 nm. But significantly, for neither spin pair are the longer substrate-induced shifts that were seen in Fig. 3d observed in the reconstituted system. Thus, when removed from the native outer-membrane environment, the structure of SB3 is altered and the large substrate induced movement of SB3 towards the periplasmic surface in the presence of the R14A mutation is no longer seen.

It should be noted that in our initial work on the V90R1-T188R1 spin pair, we failed to observe a substrate-dependent conformational change in SB3 using an isolated OM preparation where the preparation includes a sarkosyl treatment (18). This suggests that this detergent treatment of the outer membrane to remove inner membrane components is sufficient to alter the behavior of BtuB. These observations provide an explanation for why these changes in the conformation of SB3 have not been previously observed.

### Mutating either R14, D316, or both, alters SB3 conformations and populates a state where SB3 is moved towards the periplasmic surface

In earlier work, we demonstrated that breaking the ionic lock between D316 and R14 altered a conformational equilibrium in the Ton box and promoted its unfolded state (25). In this work, the effect of the R14A mutation on the Ton box equilibrium was comparable to that of a D316A mutation, but slightly enhanced for the dual R14A/D316A mutant.

Figure 5 shows a result of mutating one or both of R14 and D316 on the substrate dependent changes in SB3 as measured using the V90R1-T188R1 spin pair (the time domain data are provided in Fig. S5). In the apo state, the distributions for both the R14A and D316A mutants are similar, with peaks falling in the same positions, although the D316A mutant yields more residual area under the 3.2 nm peak than is observed for R14A. In the presence of substrate, both show significant peaks around 3.8 and 4.5 nm, and a significant peak at 2.4 nm in both the apo and vitamin B_12_ bound samples. The double R14A-D316A mutation yields a distance distribution that is more perturbed than either single mutant, with a single broad peak centered around 2.4 nm in the apo state and an increase in the longer distance components in the substrate bound state. Thus, disrupting the D316-R14 ionic lock by mutating one or both residues has a dramatic effect on the conformation of the apex of SB3 and populates a state where this binding loop has moved a significant distance towards the periplasmic interface. The data indicate that this ionic interaction plays a role in mediating allosteric changes within the BtuB core, affecting not only the Ton box equilibrium but the substrate binding loop SB3 on the extracellular surface.

**Figure 5:**
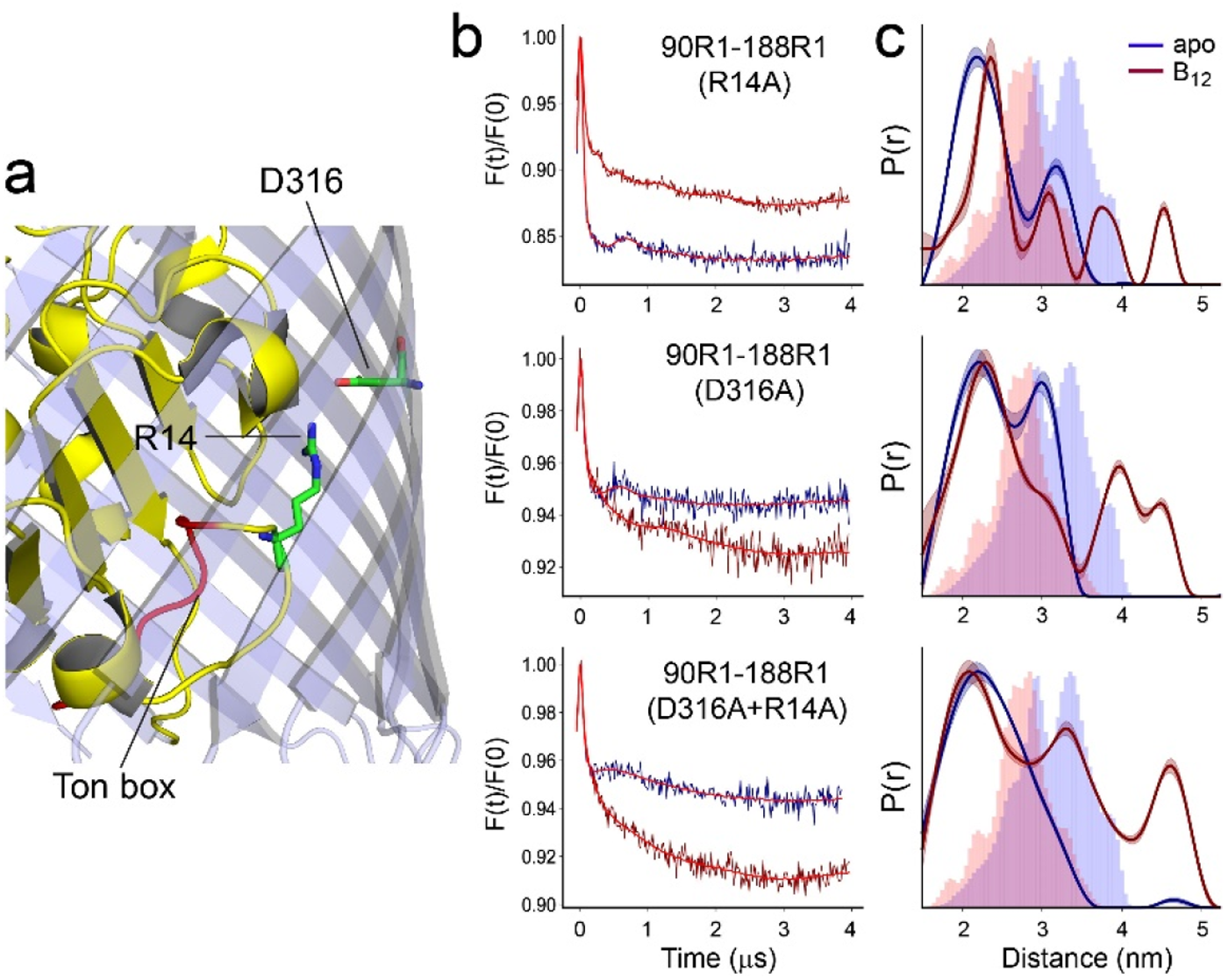
R14A, D316A or D316A-R14A have similar effects on the conformation of SB3. In (**a**) is shown the structure of BtuB highlighting the positions of the R14 and D316 side chains and the location of the Ton Box (from PDB ID: 1NQG). Background corrected DEER data are shown in (**b**) and distance distributions in (**c**) for the V90R1-T188R1 spin pair in whole cells in the presence of the R14A, D316A, or the combined R14A-D316A mutants. Data are shown for both apo (blue) and vitamin B_12_ bound (red) states. Lines through the DEER data represent the fits for the distributions shown on the right. These data were analyzed using LongDistances v932 and the model-free fitting mode. The shaded error bands in (c) represent variation due to background noise, start-time, dimensionality, and regularization. Histograms represent predicted distances generated from the *in surfo* crystal structures for PDB ID: 1NQG (light blue) and PDB ID: 1NQH (pink) using the software package MMM (21).

It should be noted that we examined the stability of the substrate-induced conformation of SB3 for times as long as 60 minutes before freezing and preparing the cells for DEER. The data are shown in Fig. S6 and indicate that the conformations are stable over time, indicating that the label is stable and not being reduced, and that conversion of the transporter back to the apo state does not occur under the conditions of the experiment.

## Discussion

The pulse EPR measurements made here in intact *Escherichia coli* indicate that substrate binding loop 3 (SB3), which includes residues 82 to 96 in the core of BtuB, undergoes a substrate-induced structural change. When the ionic interaction between R14 in the core and D316 in the barrel is broken, an alternate and more dramatic structural change in SB3 occurs upon substrate binding, where SB3 is displaced approximately 2 nm towards the periplasmic surface of BtuB. Remarkably, these structural changes do not occur when the protein is removed from the native cell environment and reconstituted into a phospholipid bilayer, indicating that features in the intact cell environment modulate the energetics of the conformational states in BtuB.

Earlier experimental work provides evidence for an allosteric coupling between the substrate binding site, the R14-D316 ion pair, and the Ton box in BtuB. Measurements made by EPR in isolated OM or reconstituted phospholipid membranes demonstrated that substrate binding partially unfolded the Ton box (10, 26), and shifted the energy of the folded and unfolded Ton box states by about 2 kcal/mol (27). When the ionic interaction between R14 in the core and D316 in the barrel was broken, the Ton box was also observed to unfold and the coupling between substrate binding and the Ton box was broken (25). In addition, a connection between the Ton box and SB3 was seen by scintillation proximity assays where both the Ton box and SB3 were found to be necessary for a TonB-dependent retention of vitamin B_12_ (28).

The connection between these sites in BtuB is also suggested by computational studies. When LPS is included in MD simulations, the interaction between R14 and D316 is weakened and the energy to unfold the Ton box reduced (24). The inclusion of LPS also alters the state of SB3. In a symmetric phospholipid bilayer, SB3 assumes the more helical form, whereas in an asymmetric membrane containing LPS, SB3 assumes an extended form. These simulations are consistent with the results presented here, except that fully populating the extended form of SB3 in our whole cell measurement (state 2 in Fig. 2d) requires substrate binding. Taken together, the results indicate that interactions made by LPS with the extracellular loops of BtuB alter conformational equilibria in the protein and may provide an explanation for the differences in the behavior of BtuB when EPR measurements are made in cells versus reconstituted phospholipid bilayers. It should be noted that the interconversion between helical and extended forms for SB3 was also absent or diminished when the V90R1-T188R1 spin pair was examined by EPR in an isolated OM preparation (18); as a result, the procedure to produce this OM preparation, which includes the use of sarkosyl, is apparently sufficient to modify the behavior of the protein.

An unexpected observation made here is that mutation of the R14-D316 ion pair alters the structure of the SB3 loop (Figs. 3d and Fig. 5). In the apo state of BtuB, this mutation enhances the extended form of SB3 (state 2 in Fig. 2b), and upon substrate binding an alternate conformational state is generated where sites 90 and 93 are extended as much as 4.5 nm from site 188 on the 3/4 extracellular loop. Among the core sites examined, this more dramatic structural change involves only SB3 as labels in the first and second substrate binding loops (SB1 and SB2) at sites 63, 65 and 72 do not exhibit any significant structural changes. Such a large structural change involving SB3 has not been previously observed, and it appears to be localized to the C-terminal side of the core.

A high-resolution model obtained by crystallography for a fragment of TonB in complex with BtuB (6) shows TonB interacting with the Ton box in an edge-to-edge manner (Fig. 6a). Since the core is largely unaltered by TonB binding, models for transport have focused on the idea that TonB alters the core structure by exerting a mechanical force on the Ton box. In particular, TonB has been proposed to function by pulling on the Ton box, which then results in an unfolding of the N-terminal region of the core (12). Single molecule pulling experiments (12) as well as steered MD simulations (11) suggest that pulling the Ton box will extract the N-terminal side of the core and eventually open a pore sufficient to allow substrate to pass. One difficulty with this model, is that both experimental and computational approaches indicate that the extraction of an extended polypeptide chain longer than the width of the periplasm is required to open this pore.

**Figure 6:**
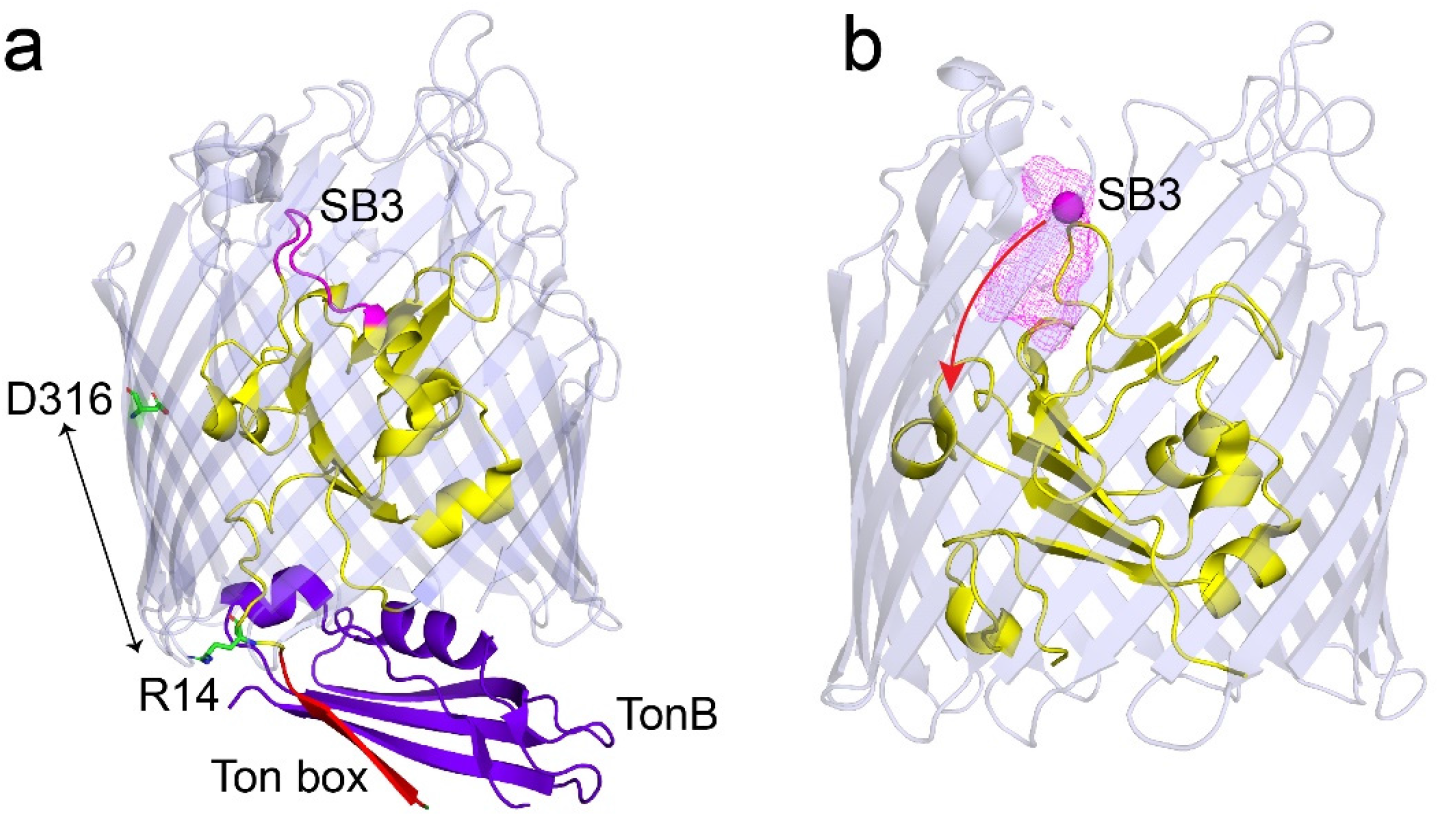
Conformational shifts in SB3 with release of R14-D316 ionic lock. (**a**) View of BtuB (PDB ID: 2GSK) in complex with the C-terminal domain of TonB (purple) showing the core (yellow) and barrel (light blue) with the substrate binding loop SB3 (magenta), and the Ton box (red), where the R14-D316 ion pair has been broken. (**b**) An Xplor-NIH simulation showing positions of the Cβ carbon (magenta mesh) on site 90 in SB3 for 300 structures that are consistent with the distributions obtained by DEER for V90R1-T188R1-R14A and V90R1-S237R1-R14A in the presence of vitamin B_12_ (see Methods). TonB binding extracts the Ton box from the core to create an edge-to-edge interaction with TonB, thereby breaking the R14-D316 ionic interaction. Breaking this interaction promotes the movement of SB3 towards the periplasmic interface of the transporter (red arrow) and may facilitate passage of vitamin B_12_.

We do not observe any significant structural changes in the N-terminal side of the core in cells, suggesting that the N-terminus may not move during transport. Rather, a significant substrate-induced structural change is found to take place in SB3 on the C-terminal side of the core when the D316-R14 ionic lock is broken. At present, we have limited restraints to generate a model and we do not know the positions of many segments in the core, but a movement in SB3 that satisfies the EPR restraints is shown in Fig. 6b. In this model, substrate binding moves SB3 into the barrel and towards the periplasm. Although we do not presently know the location of substrate under these conditions, the movement of SB3 may also drive the movement of substrate. It is interesting to note that the C-terminal side of the core has a lower side chain density than the N-terminal side, suggesting that there is more space for structural rearrangements within this region.

The structural change observed here suggests that transport does not require a mechanical pulling or rotation by TonB. When BtuB is complexed with TonB (Fig. 6a), the Ton box must disengage from the core of BtuB and extend into the periplasm (6). TonB binding exposes R14 to the periplasm, breaks the R14-D316 ionic lock and eliminates electrostatic interactions of R14 with the barrel. Thus, breaking the ionic lock through mutation of R14, as we have done here, should mimic the TonB bound state. Although the precise sequence of steps that take place during transport is not known, both substate and TonB are expected to be bound to BtuB at some point during transport. As a result, we expect that the large substrate-induced structural change observed here on the C-terminal side of the core (Fig. 6b) will mimic the state when both substrate and TonB are bound to BtuB within the cell. We conclude that TonB binding alone may promote a rearrangement in the core without the need for any mechanical pulling or rotation. Ionic interactions between the barrel and core are conserved across members of the TBDT family (8) and other transporters in this family may exhibit similar changes upon TonB binding.

The data presented here lead to an alternate model for transport where TonB binding alone initiates a structural rearrangement within the core domain of BtuB. Although more work is required to establish the steps in transport and generate a complete molecular model, this structural change may be sufficient to move the substrate into the periplasm. The energy to disrupt core-barrel electrostatic interactions and drive this structural change is provided by the free energy of binding of TonB to BtuB, which is significant and characterized by a Kd in the nM range (29). To complete the transport cycle, TonB must be disengaged from BtuB and the inner membrane complex of ExbB and ExbD may perform this function and restore the apo state. The C-terminal region of TonB is well known to form a dimer (30) and there is evidence for TonB dimers *in vivo* (31). Moreover, a TonB fragment has been observed to convert from a dimer to a monomer upon binding BtuB (29). Conceivably, TonB might be dissociated from BtuB by exchanging the strand-to-strand TonB-Ton box interaction for a strand-to-strand interaction within the TonB dimer (29, 32). In this model, the energy to drive this strand-to-strand exchange and/or regenerate the TonB monomer would be provided by the inner membrane ExbB-ExbD complex.

In summary, EPR spectroscopy in whole cells provides evidence for a substrate-induced structural transition in BtuB involving SB3 on extracellular apex of the core. The introduction of a mutation to break the R14-D316 ionic lock acting between the core and the barrel of BtuB produces an alternate structural state upon substrate addition so that SB3 is displaced as much as 2 nm into the barrel towards the periplasmic side of the protein. Under these same conditions, no movement of the N-terminal side of the core is detected. This ionic lock will be broken upon TonB binding and mutating this ionic lock is expected to mimic the TonB-bound state. As a result, this substrate-induced structural transition likely represents an intermediate in the transport process. Moreover, if TonB binding alone can significantly alter the core of BtuB, there is no longer a need to propose that TonB acts through a mechanical pulling or rotation mechanism. Remarkably, when BtuB is reconstituted into a phospholipid bilayer, these structural changes in SB3 are no longer observed, indicating that features in the native OM environment, such as the LPS, are required to populate conformational states that are important for BtuB function.

## Materials and Methods

### Cell lines and Mutants

pAG1 plasmid with WT *btuB* gene was kindly provided by late professor R. Kadner, University of Virginia. The *E. coli dsbA* null (*dsbA*^*-*^) mutant strain, RI90 (*araD139* Δ(*araABC-leu*)7679 *galU galK* Δ(*lac*)*X74 rpsL thi phoR* Δ*ara714 leu+, dsbA:: Kanr*) were obtained from the Coli Genetic Stock Center (Yale University, New Haven, CT). L63C, S65C, N72C and S93C *btuB* mutants were custom produced by Applied Biological Materials Inc. (Richmond, BC, Canada). The *btuB* mutants (L63C-T188C, S65C-T188C, N72C-T188C, V90C-T188C, V90C-S237C, S93C-T188C) with and without the R14A mutation, and V90C, V90C-T188C-D316A and V90C-T188C-R14A-D316A were engineered using polymerase chain reaction (PCR) mutagenesis. The plasmids were confirmed by sequencing and were transformed into *dsbA*^*-*^ cells. Glycerol stocks were prepared and stored at −80°C.

### Whole cells sample Preparation

*dsbA*^*-*^ cells expressing V90C-T188C, L63C, S65C, N72C, V90C and S93C BtuB were grown in minimal media (MM) supplemented with 200 µg/ml ampicillin, 0.2 % w/v glucose, 150 µM thiamine, 3 mM MgSO_4_, 300 µM CaCl_2_, 0.01 % w/v methionine and 0.01% w/v arginine (18). Cells expressing BtuB with the V90C-T188C mutation was spin labeled as described (18) and the aliquots of processed cell pellets were mixed with vitamin B_12_ (0, 1, 5, 20, 30, 60 and 100 µM final concentrations). The cells expressing L63C, S65C, N72C, V90C and S93C BtuB mutants were processed as described in Nyenhuis et al. (33).

Glycerol stocks of *dsbA*^*-*^ cells expressing L63C-T188C, S65C-T188C, N72C-T188C, V90C-T188C, S93C-T188C and V90C-S237C BtuB with and without R14A, V90C-T188C-D316A and V90C-T188C-R14A-D316A BtuB were used to directly inoculate the pre-precultures (Luria Bertani media with 200 µg/ml ampicillin), grown for 8 hours at 37°C and used to inoculate the MM precultures. The main MM cultures were inoculated with precultures, grown until OD_600_ ∼0.3 and spin labeled (33). Briefly, the cells were spin labeled with methanethiosulfonate spin label (MTSSL) ((1-oxy-2,2,5,5-tetramethylpyrrolinyl-3-methyl)methanethiosulfonate), (Cayman Chemical, Ann Arbor Michigan) in 100 mM HEPES buffer (pH 7.0) containing 2.5% (w/v) glucose with the final concentration of 7.5 nmol/ ml of cell culture at OD600 0.3 for 30 min at room temperature (RT). Spin labeled cells were washed by resuspending in 2.5% (w/v) glucose supplemented 100 mM HEPES buffers, first, at pH 7.0 and then, at pD 7.0. During the washing steps, cysteine double mutants without R14A were incubated for 15 min, while 2-5 min for mutants with R14A and 10 min for D316A and for R14A-D316A mutants. Aliquots of processed V90C-T188C-R14A samples were incubated with vitamin B_12_ for 0, 30 and 60 min at RT.

### Reconstituted BtuB sample preparation

MM main cultures of V90C-T188C and S93C-T188C with and without R14A were grown for 8 hours at 37°C. The harvested cells were used to isolate intact OM (33). After the second spin at 118,370 × g for 60 min at 4°C the pellets were resuspended in 5 mL of HEPES buffer. The OMs were solubilized in 100 mM Tris pH 8.0 buffer with 10 mM EDTA and 0.5 g of octylglucoside (OG). The OM suspension was incubated at 37°C for 10 min and 2 hours at RT and then spun at 64157 × g, 60 min at 4°C. The supernatants were used to spin label BtuB with 12 mM MTSSL at RT, overnight. BtuB was purified using 6 column volumes (CV) of wash buffer (17 mM OG, 25 mM Tris pH 8.0), 12 CV of 0-100 % gradient of elution buffer (1 M NaCl, 17 mM OG, 25 mM Tris pH 8.0) and 6 CV of 100 % elution buffer using a Q column and fractions containing BtuB were pooled. 1-palmitoyl-2-oleoyl-glycero-3-phosphocholine (POPC) (Avanti Polar Lipids, Alabaster, AL) (20 mg/ mL) was sonicated in reconstitution buffer (150 mM NaCl, 100 nM EDTA, 10 mM HEPES pH 6.5) with OG (100 mg/ mL) until clear, next, 1 ml from micelles mixture was added to each pooled BtuB mutant and incubated at RT, 40 min. BtuB was reconstituted into POPC by dialyzing OG over six buffer exchanges using reconstitution buffer and bio-beads (with minimum of six hrs dialysis per exchange). Reconstituted BtuB was pelleted by centrifugation at 23425 × g for 40 min at 4° C, resuspended in 200 μL of reconstituted buffer and further concentrated to 50 μL by using Beckman airfuge. The samples were frozen and stored at −80° C.

### EPR spectroscopy

For CW EPR, 6 uL of cell pellet, or 6 uL of cell pellet with 100 uM vitamin B_12_ were loaded into glass capillaries (0.84 O.D., VitroCom, Mountain Lakes, NJ). Capillaries were loaded into an ER 4123D dielectric resonator mounted to a Bruker EMX spectrometer. Data were taken at X-band and room temperature with 100 G sweep width, 1 G modulation, and 2 mW of incident microwave power. For pulse EPR, 16 uL of cell pellet, 20% glycerol, and 100 uM CNCbl when applicable, were combined and loaded into quartz capillaries (1.6 mm O.D., VitroCom). Samples were flash frozen in liquid nitrogen and loaded into an EN5107D2 resonator (Bruker BioSpin, Billerica, MA). Data were collected on a Bruker E580 at Q-band and 50 K using a 300 W TWT Amplifier (Applied Systems Engineering, Benbrook, TX). The dead-time free 4-pulse DEER sequence was used for all experiments, with rectangular pulses of typical lengths π/2 =10 ns and π = 20 ns, and a 75 MHz frequency separation.

### Data Processing

CW EPR spectra were normalized by dividing by the spectral second integral using in-house python scripts. Pulse EPR data were processed using LongDistances v 932 (Christian Altenbach, UCLA). Data were fit to a variable dimension background, after which the model-free mode was used for distance fitting. The value of the smoothing parameter was selected based on the L-curve, ensuring that the selected value passed through the major oscillations present in the data. Error analysis used 100 variations at the default values for background noise, start time, dimensionality, and regularization error. For data in Fig. 1D only, data were instead fit using a model-based mode with 2 Gaussian components with free position, width, and amplitude to investigate the dosage dependence of the substrate dependent shift towards a shorter distance component for the 90-188 pair. All EPR figures were generated using python scripts and the matplotlib plotting library. Protein structure images were generated using Pymol (34). Simulated distance distributions were generated using the software package MMM version 2018.2 and the default rotamer library (21, 35).

### Modeling

The 90-188, and 90-237 distributions with the 14A mutation and in the presence of vitamin B_12_ were used in the generation of a model (Fig. 6b) for the motion of the apical hatch loop using the software package Xplor-NIH (version 3.2). The starting structure for modeling was the *in surfo* structure crystal structure in the presence of cobalamin (PDB ID: 1NQH), which we previously determined to be closest to the native state of the extracellular loops in the native environment (19). In that work, we found minimal evidence for motion of the extracellular loops, and we assumed that motion was localized to the SB3 loop. The template structure was labeled *in-silico* with the R1 sidechain at the 90, 188, and 237 positions using the software package MMM (version 2017.2) and the default rotamer library. The top three rotamers were selected from the *in-silico* labeling and used to generate three input structures for ensemble calculations, with the relative weights of the three rotamers conserved from the *in-silico* labeling calculation.

During the calculation, the reference R1 sites in the barrel (188, and 237) were held entirely fixed. The R1 sidechain at site 90, the underlying SB3 loop element comprising residues 81-104, and the adjoining, unstructured hatch region comprising residues 112-124 were fully mobile during runs, while all remaining residues had fixed backbone atoms and mobile sidechains. The standard Xplor potentials BOND, ANGL, and IMPR were used in conjunction with the torsionDB and repel potentials for all elements, and DEER restraints were encoded as square well potentials using the noePot potential term. All potential terms were ensemble averaged across the three input structures. The peak positions used in the modeling were the peak centered at 4.5 nm for 90-188, and the peak at 2.8 nm for 90-237. Full peak widths were used, with the square well stopping at 5% of maximum intensity. Randomization was introduced to the calculations using the randomizeTorsions function on the starting sidechain positions of the mobile hatch elements. Following this, ten rounds of 400 step Powell minimization using all potentials and with the mobile hatch elements were used to satisfy the experimental restraints. Structure calculation was repeated until both restraints were within error.

## Acknowledgements

This work was supported by grants from the NIH (NIGMS, GM035215, S10OD025149) to DSC.

## Data availability

Raw unprocessed DEER data is available on request from D.S.C.

