## Supporting Information for "Structural intermediates observed only in intact *Escherichia coli* indicate a mechanism for TonB-dependent transport"

**This file includes:**

Figures S1 to S6
SI References

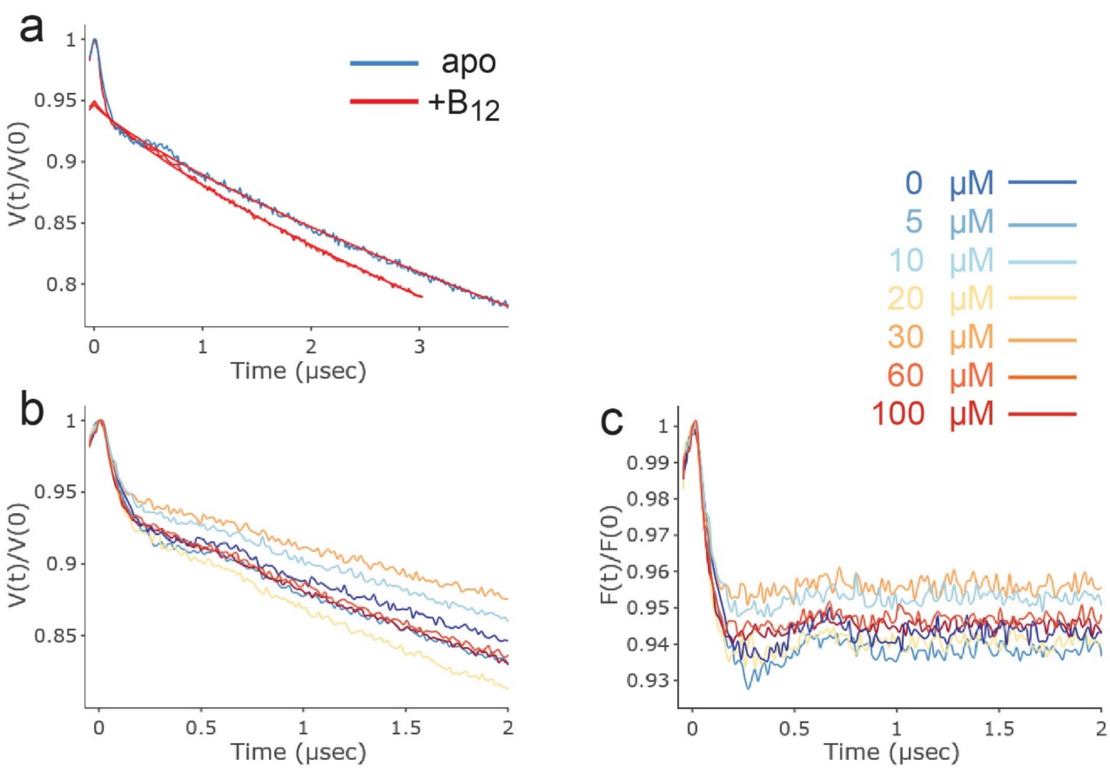


**Figure S1: Raw and background corrected DEER data for V90R1-T188R1.** In (**a**) raw DEER data, V(t)/V(0), along with the background form factor that was subtracted (straight red lines). In (**b**) raw DEER data as a function of substrate (vitamin $B_{12}$) concentration (see inset). In (c) background corrected DEER data, (F(t)/F(0), as a function of vitamin $B_{12}$ concentration (see inset). The resulting distributions are shown in Figure 2 (main text).

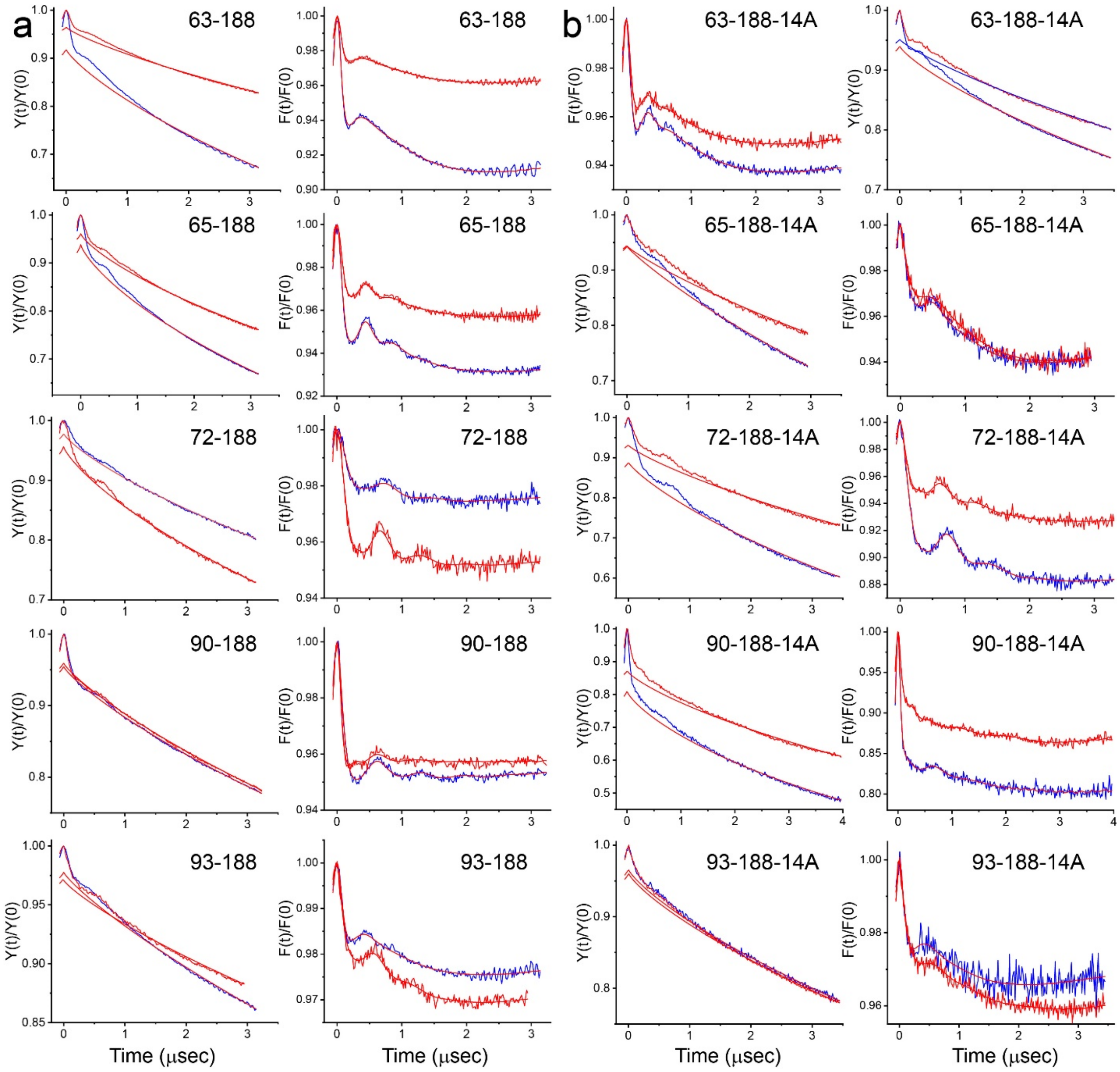


**Figure S2: Raw and background corrected DEER data for loop-core spin pairs.** In (**a**) data obtained without the R14A mutation and (**b**) data with the R14A mutation. Data for the apo state is shown in blue and the data for the vitamin $B_{12}$ bound state is shown in red. For each of the 20 sets of distributions shown in Figure 3 (main text) are shown the raw DEER data, V(t)/V(0) (left panels), along with the background form factor that was subtracted (straight red lines). The background corrected DEER data, (F(t)/F(0) (right panels), are shown along with the fits to the data (straight red lines) used to generate the distributions in Figure 3 (main text).

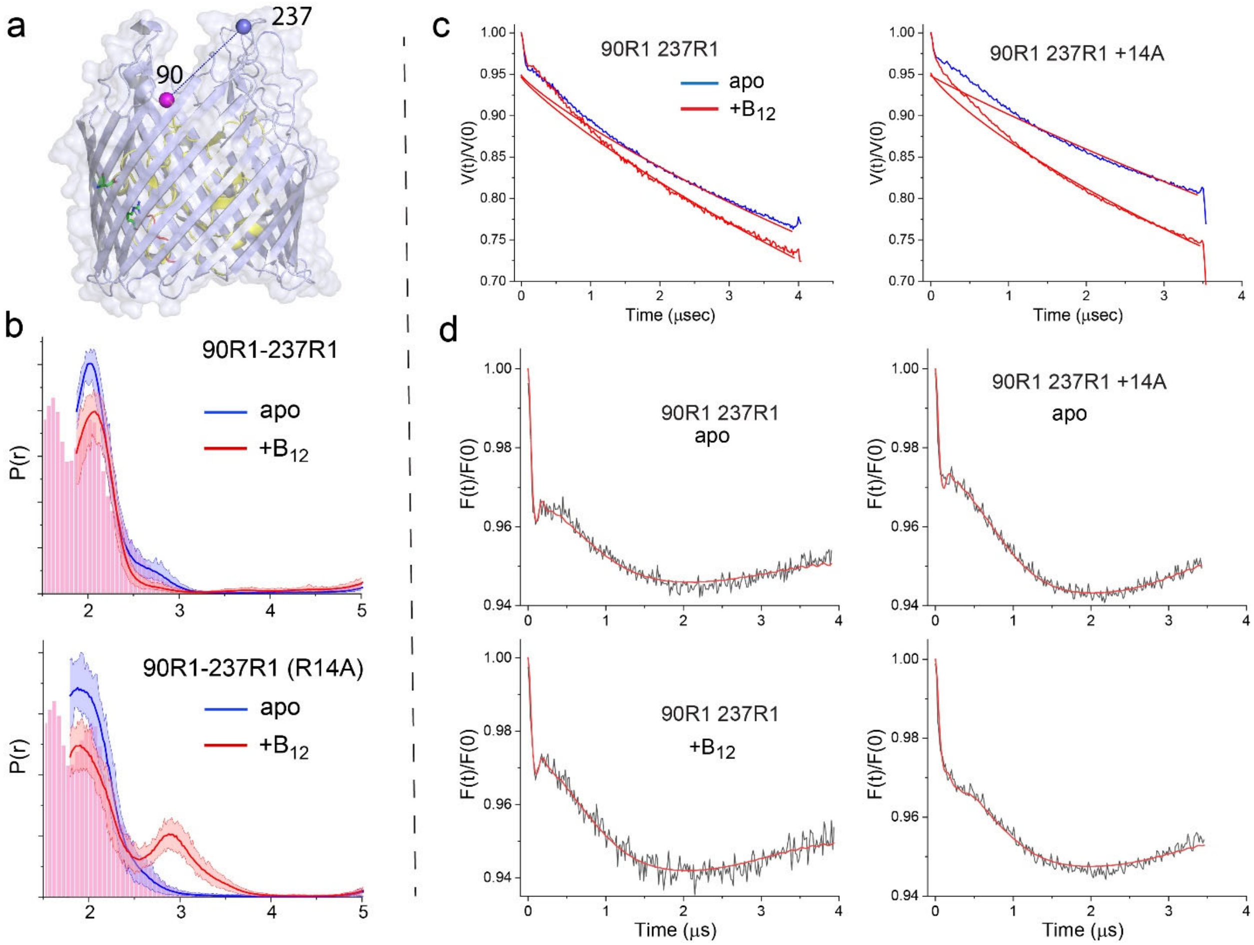


**Figure S3. The substrate-dependent movement in SB3 promoted by the R14A mutation is observed at site 237.** Shown in (**a**) is a side view of sites 90 and 237 (PDB ID:1NQH). In (**b**) is shown the distribution for the V90R1-S237R1 spin pair in the absence and presence of R14A with and without vitamin $B_{12}$, where the histrogram represents prediction from the Vitamin B12 bound structure (PDB ID: 1NHQ). A significant portion of the predicted distance distribution is below the minimum range observable using DEER (1.5-2 nm). Shown in (**c**) are the raw data and background for the V90R1-S237R1 spin pair without and with the R14A mutation, and in (d) are the background correct DEER data. These data were analyzed using the DeerNet routine (1) in DEERAnalysis (2). Histograms are predicted distances generated from the *in surfo* crystal structures (PDB ID: 1NQG (blue) and PDB ID: 1NQH (red)) using MMM (3).

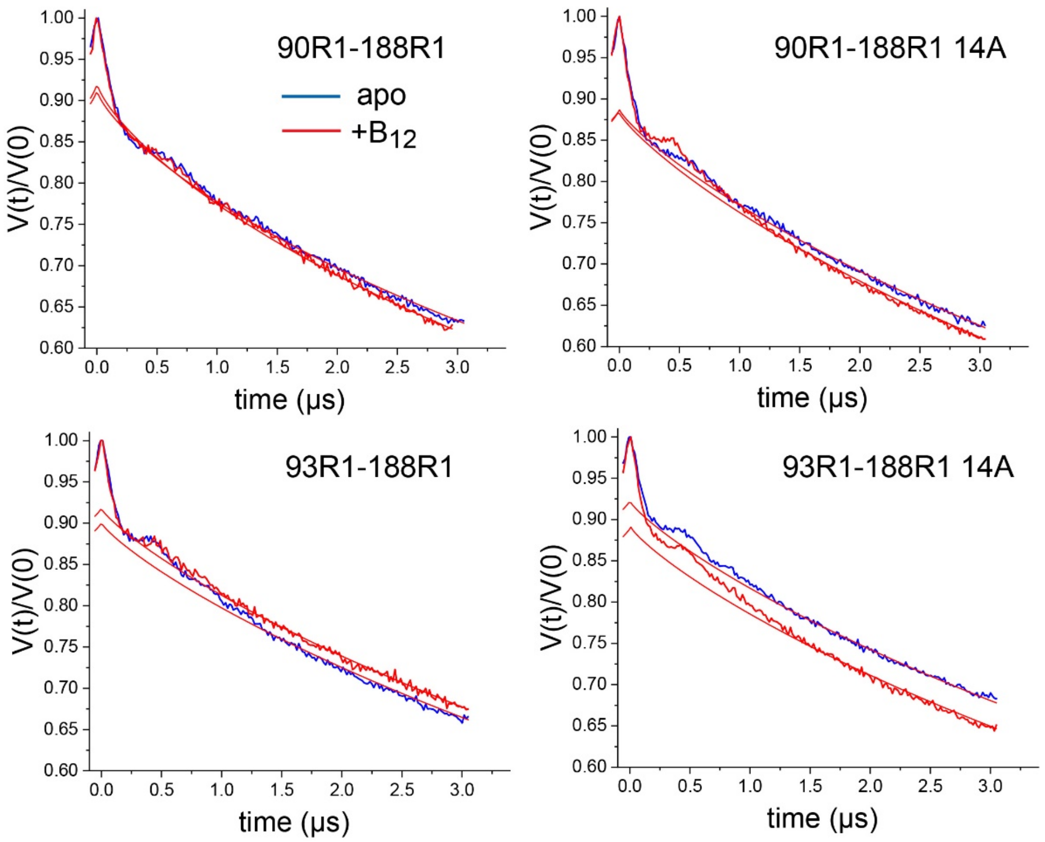


**Figure S4. Raw DEER data for the V90R1-T188R1 and S93R1-T188R1 spin pairs purified and reconstituted into POPC.** Solid red lines indicate the background form factors that were used to obtain the background corrected data shown in Figure 5 (main text).

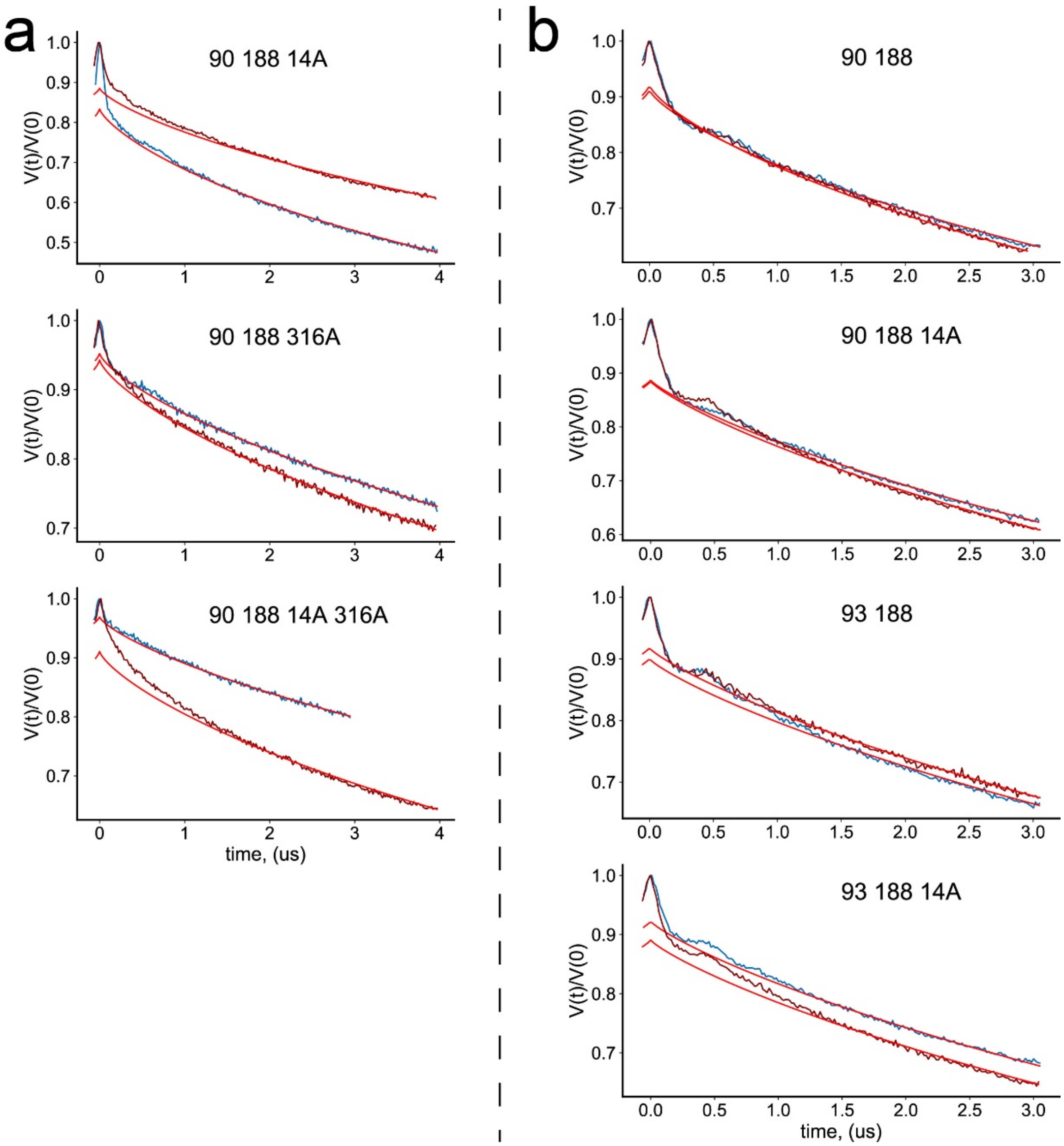


**Figure S5: Raw DEER data for the V90R1-T188R1 spin pair in the presence of the R14A, D316A and R14A/D316A mutations.** Solid red lines indicate the background form factors that were used to obtain the background corrected data shown in Figure 6 (main text).

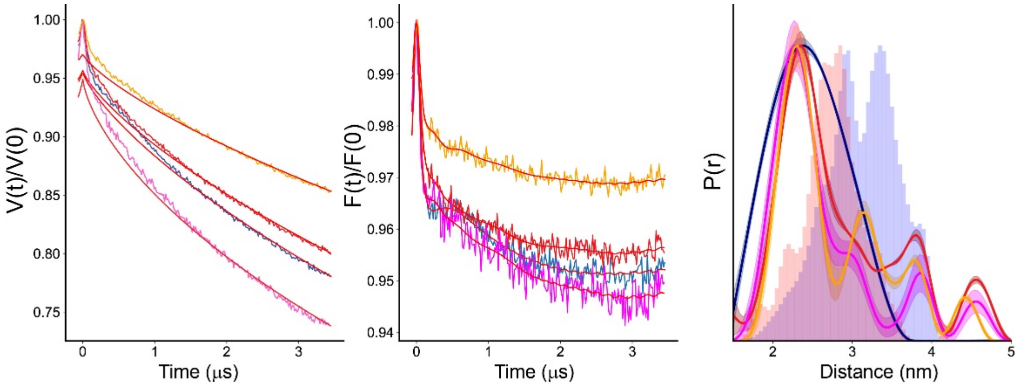


**Figure S6: Time domain (left), dipolar (middle) and distance (right) data obtained for the V90R1-T188R1 pair with the R14A mutation at various time points show relative stability of the new distances.** The initial apo (blue) and vitamin $B_{12}$ (red) traces were processed and frozen more quickly than in other figures, and it can be seen that the starting distribution, particularly for the apo condition, is broader with a single peak around 2.6 nm. From 0 to 30 minutes (magenta) after addition of substrate, there is a narrowing of the distance elements, which continues to 60 minutes after addition (orange). Despite this, and the loss of signal by 60 minutes seen in the decrease in modulation depth, the locations of the peaks are not altered, indicating that either these conformations, or their percentage abundance, are stable on the 60-minute time scale. Data were analyzed using LongDistances v932 and the model-free fitting regime. Histograms are predicted distances generated from the *in surfo* crystal structures 1NQG and 1NQH using the software package MMM.
